# Localization of *Kif1c* mRNA to cell protrusions dictates binding partner specificity of the encoded protein

**DOI:** 10.1101/2022.11.07.515531

**Authors:** Megan L. Norris, Joshua T. Mendell

## Abstract

Subcellular localization of messenger RNA (mRNA) is a widespread phenomenon that can impact the regulation and function of the encoded protein. In non-neuronal cells, a subset of mRNAs localize to cell protrusions and proper mRNA localization is required for cell migration. However, the mechanisms by which mRNA localization regulates protein function in this setting remain unclear. Here, we examined the functional consequences of localization of the mRNA encoding KIF1C. KIF1C is a kinesin motor protein required for cell migration and mRNA trafficking, including trafficking of its own mRNA. We show that *Kif1c* mRNA localization does not regulate KIF1C protein abundance, distribution, or ability to traffic other mRNAs. Conversely, robust *Kif1c* mRNA localization is required for directed cell migration. We used mass spectrometry to identify binding partners of endogenous KIF1C, which revealed dramatic dysregulation of the number and identity of KIF1C interactors in response to *Kif1c* mRNA mis-localization. These results therefore uncovered a mechanistic connection between mRNA localization to cell protrusions and the specificity of protein-protein interactions. We anticipate that this mechanism is not limited to *Kif1c* and is likely to be a general principle used by protrusion-enriched mRNAs in non-neuronal cells.

## INTRODUCTION

Subcellular localization of mRNA is a widely-occurring form of post-transcriptional regulation that can tune protein output in space and time. While mRNA localization has been most extensively studied in large, highly asymmetrical systems, such as oocytes and neurons, it also occurs in smaller cells including both mesenchymal and epithelial cell types (reviewed in ^1,2^). In non-neuronal migratory cells, disruption of mRNA localization to cellular protrusions routinely leads to defective cell migration and has been shown to affect processes *in vivo* including cancer cell invasion and blood vessel morphogenesis^3–6^. It was predicted that mechanistic principles of mRNA localization in non-neuronal cells would be similar to those of neurons, but recent work has challenged that assumption. In neuronal systems, *cis*-elements in localized mRNAs are bound by RNA binding proteins, which are then carried by molecular motors into neurites and docked using the same or different proteins^7^. The mRNAs are generally translationally silent during transport and undergo local translation at their final site in the neurite, often in response to a stimulus. In contrast, global mRNA and protein localization patterns are not correlated in protrusions of non-neuronal cells^8^. Furthermore, at least some localized mRNAs are translated en route to non-neuronal cell protrusions, only to be translationally silenced at their destination^9^. Thus, the functional impact of mRNA localization to protrusions in non-neuronal cells cannot be directly inferred from principles established in neurons, necessitating further empirical characterization of the molecular and phenotypic consequences of mRNA mis-localization in non-neuronal cell types.

Work to date has revealed at least two classes of protrusion-localized RNAs in non-neuronal cells: those that require the tumor suppressor APC to localize (“APC-dependent”), and those that do not (“APC-independent”)^4,10^. While important aspects of the localization mechanisms for both groups remain to be elucidated, key insights support the hypothesis that each group uses its own set of general principles for localization. For example, multiple APC-dependent mRNAs have been definitively shown or predicted to have guanine and adenine (GA) rich *cis*-elements that mark them for trafficking^3,5,6,11^. Many, if not all, APC-dependent mRNAs also require the kinesin KIF1C for trafficking^12^. Conversely, many APC-independent mRNAs, which includes a large number of ribosomal protein mRNAs, use LARP family members to localize^4,13,14^. Intriguingly, these two classes of localized mRNAs seem to populate different types or stages of protrusions^4^. A glimpse into the molecular consequences of localizing APC-dependent mRNAs has been provided by detailed studies of *Rab13*. Localization of *Rab13* mRNA to protrusions does not affect RAB13 protein abundance or distribution, but regulates co-translational loading of a key interacting partner^6^. Without this regulated loading, cells with mis-localized *Rab13* mRNA phenocopy RAB13 knockdown cells, exhibiting a migration defect. However, whether mRNA localization to protrusions commonly regulates protein-protein interactions, or whether this mechanism is unique to *Rab13*, remains to be determined.

Another representative APC-dependent mRNA is *Kif1c*, which encodes a kinesin-3 motor protein^15^. *Kif1c* is one of the most commonly localized mRNAs across cell types and has been observed in protrusions in diverse cell lines including HeLa, mouse fibroblasts (NIH3T3), and human umbilical vein endothelial cells (HUVECs)^4,5,16^. Previous work has determined that the *Kif1c cis*-localization element is sufficient to drive localization of reporter mRNAs and is predicted to include GA elements, similar to other APC-dependent mRNAs^3,5,6^. KIF1C protein is required for directed cell migration and proper transport of α5β1 integrin to focal adhesions, specifically in the rear of the cell^17^. As described above, KIF1C also interacts with APC to traffic its own mRNA, as well as other APC-dependent mRNAs^12^. These studies paint a picture in which *Kif1c* mRNA localization is a carefully orchestrated process shared across organisms and cell types and predicts that *Kif1c* mRNA localization and protein function may be intertwined. Nevertheless, it is unclear if localization of the *Kif1c* mRNA is required for proper KIF1C-mediated mRNA trafficking or cell migration.

Here, we tease apart the multiple functions of KIF1C and uncover their specific dependencies on mRNA localization. We find that *Kif1c* mRNA localization is dispensable for trafficking other APC-dependent mRNAs and has no effect on KIF1C protein abundance or distribution. Conversely, mis-localization of *Kif1c* mRNA leads to a broad reprogramming of KIF1C protein-protein interactions and results in defective cell migration. Thus, *Kif1c* mRNA localization appears to be critical for the establishment of distinct functional pools of KIF1C protein that are produced in different cellular compartments, thereby providing access to distinct sets of cargoes.

## RESULTS

### Identification of Kif1c as a model localized RNA

To gain mechanistic insight into the manner by which mRNA localization to protrusions affects cellular behavior, we focused on the mouse melanoma cell line YUMM1.7. This cell line is phenotypically rich, amenable to genetic manipulation, and genetically engineered to resemble human melanoma through activated *Braf* and deactivated *Pten* and *Cdkna*^18^ (Figure 1A). To identify RNAs that are localized to protrusions in these melanoma cells, we carried out a candidate prioritization pipeline that consists of four phases: identification, comparative analysis, phenotypic screening, and cursory *cis*-element identification.

**Figure 1.**
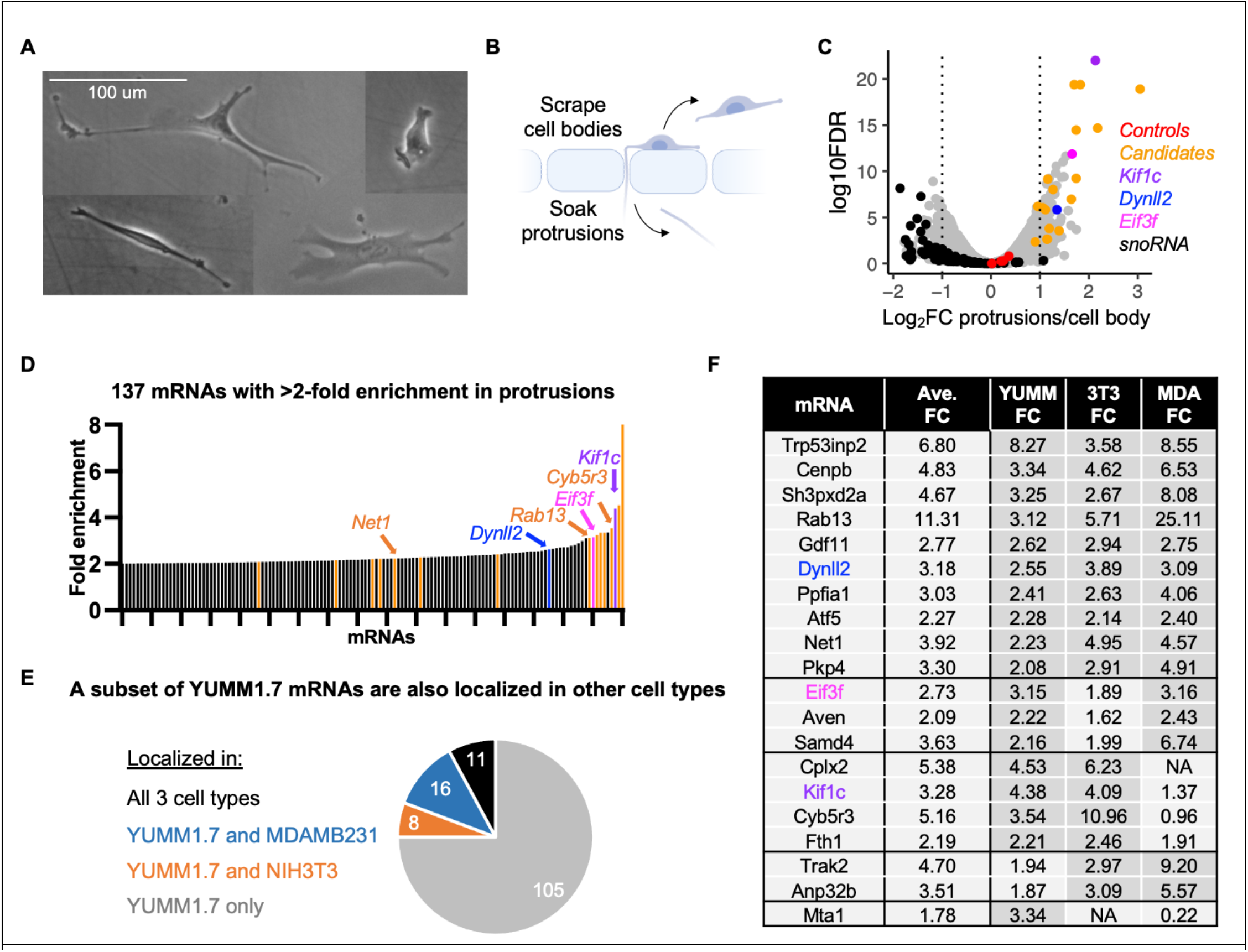
Identification and prioritization of mRNAs localized to protrusions in melanoma cells. **A**. YUMM1.7 cells are a morphologically diverse and migratory cell type. **B**. YUMM1.7 cells were fractionated into protrusions and cell bodies using microporous membranes. **C**. RNA-seq results for fractionated YUMM1.7 cells. Control mRNAs such as *Ppia, Ywhaz, RhoA* and *Arpc3* are uniformly distributed between protrusions and cell bodies (red dots), while nucleolar RNAs are depleted from protrusions (black). **D**. Protrusion enrichment values for the top 137 localized mRNAs in YUMM1.7 cells. In **C and D**, *Dynll2, Eif3f, Kif1c* and other candidate genes are denoted in color. **E**. 35 mRNAs that localize in melanoma cells are also localized in breast cancer cells (blue), embryonic fibroblasts (orange) or all three cell types (black). **F**. Candidate mRNAs and their protrusion-enrichment patterns in three non-neuronal cell lines. Table is arranged based on the candidates’ enrichments across three, two or one cell type(s).

#### Phase 1, Identification

Cells were plated on microporous membranes coated on the underside with fibronectin. Protrusions, but not cell bodies, are able to fit through the 1 μm pores (Figure 1B). Cell bodies were collected from the top of the membrane by scraping, followed by collection of protrusions by soaking the resulting membranes in lysis buffer. RNA was isolated from protrusion and cell body samples and subjected to RNA sequencing. mRNAs that were enriched by at least two-fold in the protrusion versus the cell body fraction were considered enriched (Figure 1C-D). mRNAs with well-studied subcellular localization patterns behaved as expected in these data. For example *Arpc3* and *RhoA* mRNAs were uniformly distributed (near 0 in Figure 1C, red dots), while *Rab13, Cyb5r3* and *Net1*, which are known to be localized in other non-neuronal cells, were enriched in protrusions (Figure 1D-F).

#### Phase 2, Comparative analysis

To narrow down our list of candidates, we excluded mRNAs with low expression in cell bodies (TPM < 1), resulting in 137 mRNAs that are two-fold or more enriched in protrusions. Next, we selected nine of the most enriched mRNAs and mRNAs with shared enrichment across three non-neuronal cell types (fibroblasts, breast cancer, and melanoma) (11 additional mRNAs) (Figure 1 D-F). We reasoned that, though the absolute enrichment of these 11 genes may be more moderate, they may have a higher chance of being biologically interesting due to their shared enrichment across cell types. In this way we chose 20 candidates that were particularly strongly and/or commonly enriched in protrusions (Figure 1F).

#### Phase 3, Phenotypic screening

To identify candidates whose localization was most likely to regulate an observable phenotype in YUMM1.7 cells, we initially used gene loss of function as a proxy. This strategy assumed that a gene with a detectable loss of function phenotype would be more likely to manifest a phenotype upon mis-localization of the encoded mRNA, although the severity or nature of the phenotype might be different. To do this, we individually targeted the coding sequence of each candidate gene from phase 2 using lentivirally-delivered CRISPR/Cas9 components. This resulted in pools of cells containing heterogenous mutations in the coding sequence for the targeted gene. For each gene we tested two separate guides and efficacy of targeting was determined using high-throughput amplicon sequencing. In most cases, both guides for a given gene resulted in a >80% frequency of indels at the target site, and differences in targeting efficacy were taken into consideration when analyzing phenotypes. Cell pools were also made with non-targeting guides to use as controls in all phenotypic assays. After targeting, cell pools were systematically screened for phenotypic changes in cell shape or cell motility using live-cell tracking (Figure 2A). The tracking assays were performed at two separate time points (one and two weeks post selection) to account for differences in cellular viability and/or protein stability. Thus, at the completion of the assays, each gene had been analyzed four times (two guides x two timepoints). For each gene, we selected the single treatment (guide and timepoint) with the strongest phenotype to be plotted as a representative (Figure 2B-D). We observed changes in cell length in both directions, with some cells becoming overall longer and others shorter depending on the targeted gene (Figure 2B). To quantify cell motility, we used two metrics: speed and persistence (how straight the cell migrates). Cells in which *Dynll2* or *Eif3f* had been targeted migrated the fastest (Figure 2C), and cells targeted for *Kif1c, Gdf11* and *Eif3f* migrated the least straight (Figure 2D). We selected *Kif1c* and *Eif3f* for further analysis because they have strong migration phenotypes and are amongst the top 10 most localized mRNAs in YUMM1.7 cells (Figure 1D and 1F). We also selected *Dynll2*, which has a migration speed phenotype, is among the top 20 most localized mRNAs in YUMM1.7 cells, and is more highly expressed than *Gdf11*.

**Figure 2.**
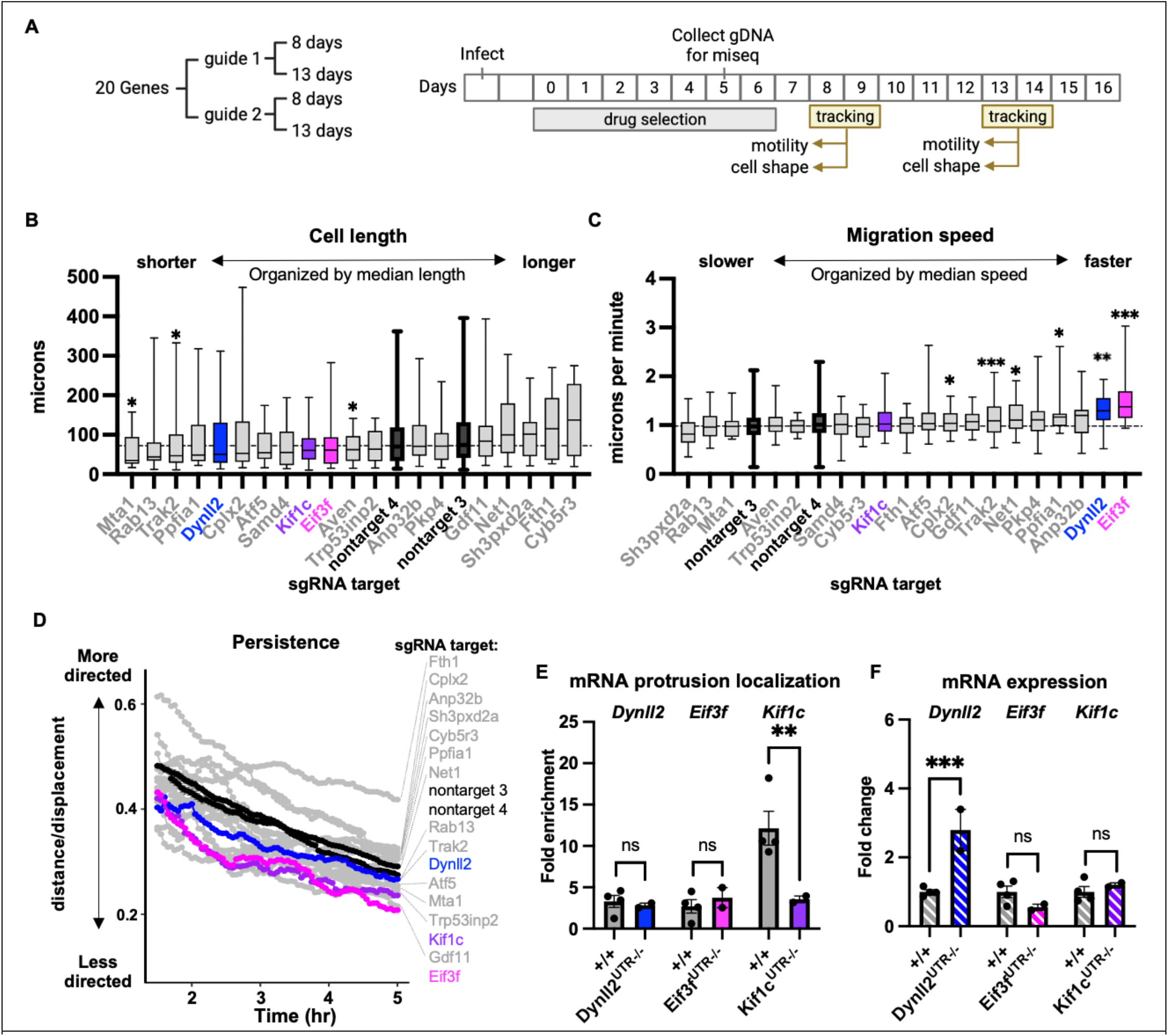
Functional prioritization of localized mRNAs and identification of *Kif1c* as a model for further study. **A**. Outline of phenotypic screen. Coding sequences of candidate genes were targeted individually with CRISPR/Cas9, selected with puromycin, then analyzed for phenotypes at two different time points. **B-D**. Only the top treatment (guide and timepoint) is shown for each candidate gene. Genes ordered based on their median measurement; horizontal dashed lines mark the median of nontarget control treatments. Nontarget guides, *Dynll2, Eif3f* and *Kif1c* are denoted in black or color. All other candidates are in gray. Cells were tracked migrating on uncoated plastic dishes for 5 hours. N=15+ cells per assay for candidate genes. N=150+ cells per assay for nontarget guides. **B**. Cell length was measured from still frames. **C**. Average cell speed (total path length/time). **D**. Migratory persistence over time. Showing mean + SEM for nontarget guides. **E-F**. qRT-PCR on cells with endogenous 3’ UTR deletions. All qRT-PCR data was normalized to *Ywhaz*, which was validated as stably expressed and uniformly distributed in all genetic contexts under study. All statistical tests are ordinary one-way anova compared to +/+. *** p<0.001; **p < 0.01; *p < 0.05.

#### *Phase 4*, cis-*element identification*

mRNA localization elements are commonly but not always found in the 3’ UTR. To test if the 3’ UTR harbored the localization element of *Dynll2, Eif3f* and *Kif1c*, we simultaneously introduced Cas9 and two sgRNAs, one targeting just after the stop codon, and one targeting just prior to the poly-adenylation sequence, and isolated clonal populations of cells with homozygous 3’ UTR deletions. To focus on the role of mRNA localization specifically, we sought to identify candidate genes whose localization elements do not also affect mRNA expression levels. Neither loss of the *Dynll2* 3’ UTR nor the *Eif3f* 3’ UTR had a significant effect on mRNA localization, suggesting that the *cis*-element is elsewhere in these mRNAs (Figure 2E), although loss of the *Dynll2* 3’ UTR did result in increased mRNA expression (Figure 2F). In contrast, loss of the *Kif1c* 3’ UTR abrogated mRNA localization without affecting mRNA expression, making it the best overall candidate for mechanistic study (Figure 2E-F).

### Kif1c mRNA localization requires a short GA-rich element in the proximal 3’ UTR

The *Kif1c* 3’ UTR is over 3,000 nucleotides long, so we next sought to identify minimal deletions that impair mRNA localization. A previous study using a reporter construct and the *Kif1c* 3’ UTR suggested that GA regions at the proximal end of the 3’ UTR may be most critical^5^. To test this, we used dual sgRNAs to delete larger (∼130nt) and smaller (∼50) portions of the GA elements in the endogenous *Kif1c* 3’ UTR (Figure 3A-B). Cellular fractionation confirmed that the GA elements are required for localization. Compared to deletion of the entire *Kif1c* 3’ UTR, deletion of most of the GA elements (*Kif1c*^Δ*GA1*,2^) or just the first GA element (*Kif1c*^Δ*GA1*^) resulted in equivalent mis-localization, while deletion of the second GA element (*Kif1c*^*ΔGA2*^) had a slightly weaker, though still significant, effect (Figure 3C). None of the deletions affected total *Kif1c* mRNA abundance or cell viability, except for a modest growth deficiency in *Kif1c*^*ΔGA1*^, which is not uncommon when generating clonal cell lines (Figure 3D-E). We conclude that *Kif1c*, like *Net1* and *Rab13*, uses GA-rich elements for protrusion localization.

**Figure 3.**
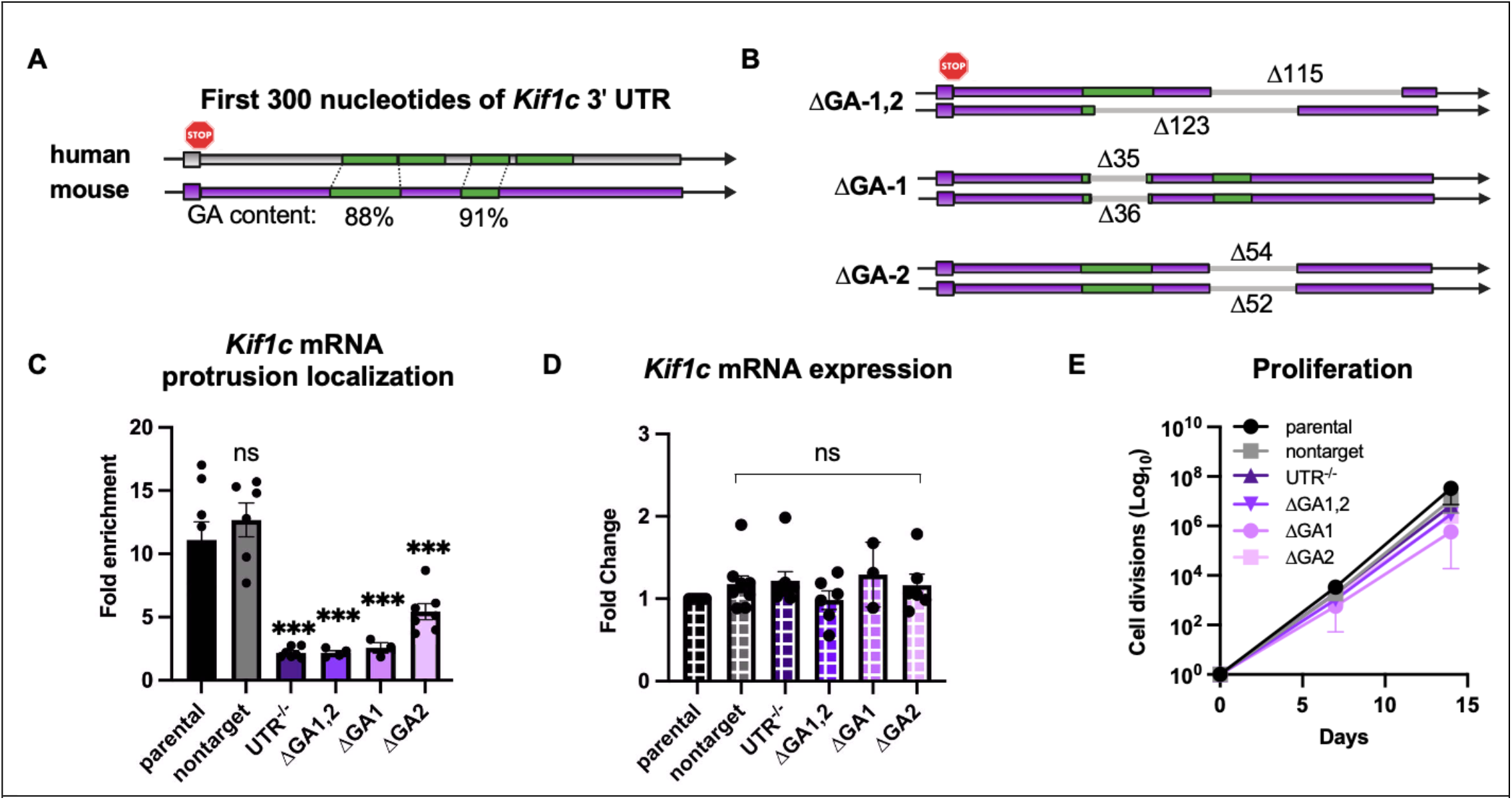
The *Kif1c* mRNA localization element is made up of multiple GA-rich sequences. **A-B**. Schematic of the first 300 nucleotides of the *Kif1c* 3’ UTR, which is 3,076 nucleotides long. GA-rich elements (green) are highly conserved between mice and humans. **B**. Schematic of endogenous deletions generated in this study. The various ΔGA clonal cell lines harbor unique deletions on either allele. **C-D**. qRT-PCR of *Kif1c* mRNA in cells with endogenous deletions. All qRT-PCR data was normalized to *Ppia* and *Ywhaz*, which were validated to be stably expressed and uniformly distributed in all genetic contexts under study. All statistical tests are ordinary one-way anova compared to parental. *** p<0.001. **E**. Cell divisions measured every 2-3 days for 2 weeks. Repeated 2-6 times per genotype.

### Kif1c mRNA localization does not regulate KIF1C protein abundance or distribution

mRNA localization can be an efficient way to regulate protein abundance and/or distribution. To test whether *Kif1c* mRNA localization affects these parameters, we first isolated protein from various *Kif1c*^*+/+*^ or *Kif1c*^*ΔGA*^ cell lines. No change in protein abundance, as measured by western blotting, was detected (Figure 4A). To visualize KIF1C spatially within the cell, we used homologous recombination to tag the C-terminus of endogenous KIF1C with mCherry (Figure 4B). Endogenous KIF1C:mCherry signal was localized perinuclearly and in cell protrusions, as previously described (Figure 4C)^17^. Next, we used CRISPR/Cas9 with dual sgRNAs to delete the GA1,2 localization element specifically on the mCherry-tagged allele. Localization of KIF1C:mCherry^ΔGA1,2^ was indistinguishable from KIF1C:mCherry^+/+^ and was readily found in cell protrusions (Figure 4D-E). Finally, we used flow cytometry to measure changes in mCherry abundance. In agreement with our western blot data, no change in protein abundance was observed (Figure 4F). Thus, *Kif1c* mRNA localization does not detectably regulate KIF1C protein abundance or distribution in the cell.

**Figure 4.**
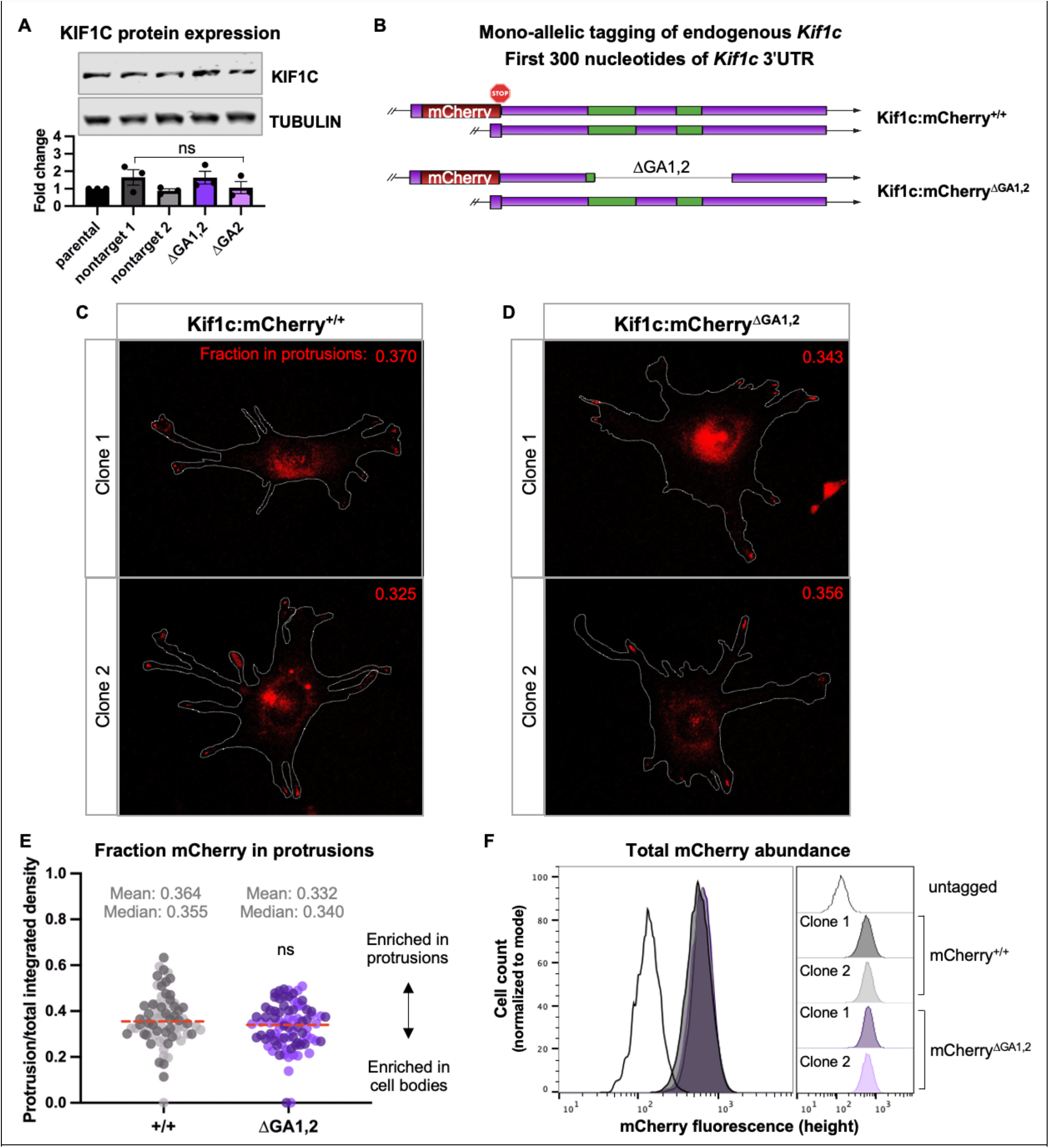
*Kif1c* mRNA localization does not affect KIF1C protein abundance or distribution. **A**. Representative western blot of total protein lysate and quantification of three independent replicates. **B**. Schematic of mCherry insertion into the *Kif1c* endogenous locus and subsequent deletion of ΔGA1,2 element. ΔGA1,2 clones 1 and 2 have 123 and 164 nucleotides deleted, respectively. **C-D**. Representative images of live cells expressing KIF1C:mCherry from localized mRNA **(C)** or mis-localized mRNA **(D)**. An outline of the cells is provided in gray. Inset number is the value shown in (E) for the imaged cell. **E**. Quantification of mCherry fluorescence present in protrusions quantified from cells represented in C and D. Separate clones for each genotype are represented as lighter and darker points. Experiment was repeated three times with 10 or more cells per clone per experiment. Red dashed line is the median. Statistical test is unpaired t test. **F**. Quantification of total cellular mCherry fluorescence using flow cytometry. >12,000 cells per population. The same data is shown as an overlay (left) and individual populations (right).

### Kif1c mRNA localization is required for directional cell migration but is dispensable for APC-dependent mRNA trafficking

Knockdown of KIF1C has been independently shown to cause defects in directed cell migration and mRNA trafficking^12,17^. Whether these processes are interdependent is unclear. Likewise, a causal relationship between *Kif1c* mRNA localization and downstream KIF1C functions has not been established. We hypothesized that mis-localization of *Kif1c* mRNA would lead to defects in KIF1C function and result in cell migration and mRNA trafficking defects. To test this, we first generated clonal loss-of-function mutants (*Kif1c*^*LOF*^) to serve as positive controls in our assays. To this end we targeted the *Kif1c* coding sequence with CRISPR/Cas9 and obtained clonal *Kif1c*^*LOF*^ cells with bi-allelic early frameshift mutations, which introduce premature stop codons. *Kif1c*^*LOF*^ mutants have decreased *Kif1c* mRNA expression and no detectable protein (Figure 5A-C).

**Figure 5.**
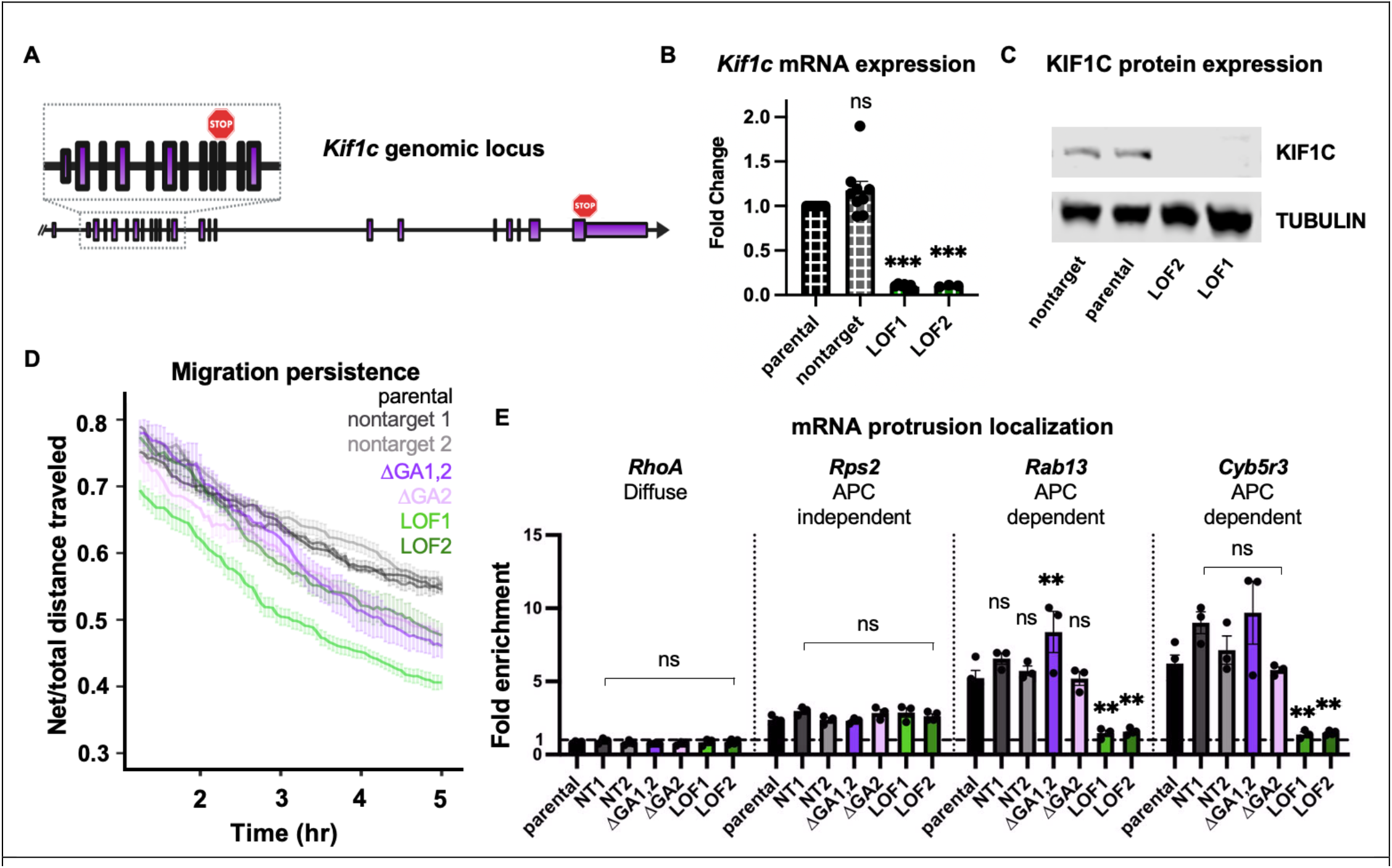
*Kif1c* mRNA localization is required for directed cell migration but dispensable for mRNA trafficking. **A**. Schematic of the *Kif1c* genomic locus. Stop sign in the inset denotes location of premature termination codon in LOF alleles. LOF1 and LOF2 harbor 2-and 4-base pair deletions in exon 11, respectively. **B**. qRT-PCR on cells with endogenous deletions. **C**. Representative western blot of total protein lysates. Experiment was repeated three times. **D**. Cells were plated sparsely on fibronectin-coated plastic dishes for 18-24 hours, then tracked every three minutes for at least 5 hours. 15 or more cells per genotype per experiment. Experiment repeated three or more times per genotype. Error bars show SEM. **E**. qRT-PCR measurement of mRNA enrichment in protrusions in the indicated cell lines. All qRT-PCR data was normalized to *Ppia* and *Ywhaz*. All statistical tests are ordinary one-way anova compared to parental. *** p<0.001.** p<0.01.

We first measured the ability of *Kif1c*^*+/+*^, *Kif1c*^*ΔGA*^, and *Kif1c*^*LOF*^ cells to undergo directed cell migration. Cells were plated sparsely on fibronectin coated plates, allowed to acclimate overnight, then tracked for 5 hours. As expected, *Kif1c*^*LOF*^ cells did not migrate as persistently as *Kif1c*^*+/+*^ cells. Consistent with our hypothesis that mRNA localization is required for KIF1C function, *Kif1c*^*ΔGA*^ cells resembled *Kif1c*^*LOF*^ cells and had diminished directed cell migration (Figure 5D). Next, we measured mRNA trafficking by fractionating cells and examining the localization of multiple protrusion-enriched mRNAs. As expected, the APC-dependent mRNAs *Rab13* and *Cyb5r3* did not localize properly in *Kif1c*^*LOF*^ cells, while the APC-independent mRNA, *Rps2*, localized normally. Surprisingly, the KIF1C-dependent mRNAs, *Rab13* and *Cyb5r3*, were properly localized in *Kif1c*^*ΔGA*^ cells (Figure 5E). Thus, in contrast with the prediction that *Kif1c* mRNA localization is required for KIF1C-dependent mRNA trafficking, we observed that KIF1C’s role in mRNA trafficking does not require *Kif1c* mRNA to itself be localized. Taken together, these results demonstrate that the functions of KIF1C in cell migration and mRNA trafficking are separable and suggest that there are multiple pools of KIF1C protein that are predicated on localization of the *Kif1c* mRNA. Furthermore, the observation that mRNA trafficking occurs properly in *Kif1c*^*ΔGA*^ cells is consistent with our observation that these cells still have KIF1C in their protrusions, the destination of KIF1C when trafficking other mRNAs (Figure 4C-E).

### Identification of endogenous KIF1C-interacting proteins

We have established that *Kif1c* mRNA localization is required for proper cell migration. However, as mRNA localization did not affect the distribution or abundance of KIF1C protein, the question remains as to how mRNA localization regulates KIF1C function. Like other kinesins, KIF1C carries cargoes along microtubules and can modulate different cellular processes dependent on which cargo it carries. For example, KIF1C interacts with APC to mediate mRNA trafficking, while KIF1C regulates directed cell migration through transport of α5β1-integrins to the rear of the cell^12,17^. We hypothesized that changes in mRNA localization may affect KIF1C function by impacting its binding partners, since mRNAs in the cell body or protrusions would be translated in the context of distinct sets of potential interactors of the encoded protein. To test this, we carried out immunoprecipitation (IP) of endogenous KIF1C from *Kif1c*^*+/+*^ and *Kif1c*^*ΔGA1,2*^ cells, with *Kif1c*^*LOF*^ cells included as a negative control, followed by mass spectrometry to identify interacting proteins. We chose to IP endogenous KIF1C because over-expression of KIF1C leads to cellular phenotypes, such as increased golgi object size^19^ and length of cell tails^17^, and we did not want to overwhelm any regulatory processes acting on the protein.

Among the interactors detected in *Kif1c*^*+/+*^, *Kif1c*^*ΔGA1,2*^, or both cell lines, we observed known KIF1C binding partners such as RAB6 and HOOK3^19,20^ (Figure 6A). A substantial number of the interactors (over 40%) were related to the cytoskeleton, integrin signaling, and/or vesicle transport (colored dots in Figure 6A). We looked for overrepresentation of biological pathways by comparing the interacting proteins to all genes expressed in YUMM1.7 cells. The broad classes of integrin signaling and cytoskeletal proteins are indeed enriched (5.8 and 6.5 fold, respectively). More striking, however, is the overrepresentation of barbed-end actin filament capping proteins (17 fold) and Arp2/3 complex proteins (26.5 fold). In other cell types, KIF1C is thought to provide an interface for actin and tubulin, which is required for the formation of actin-based invasive protrusions called podosomes. Whether KIF1C is also serving in this role in melanoma cells or is interfacing with actin to serve another role is unclear.

**Figure 6.**
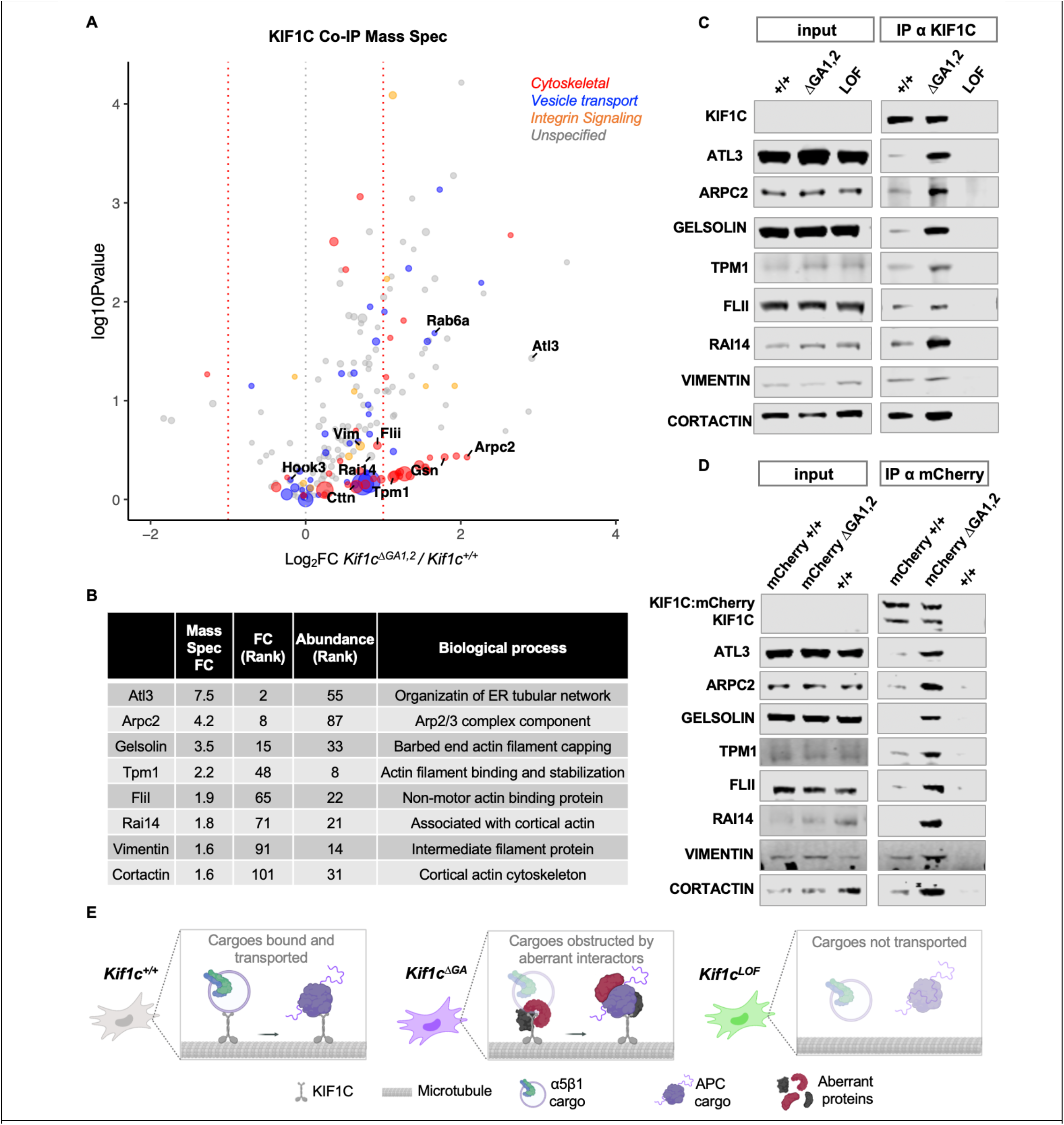
Identification of endogenous KIF1C binding partners and their regulation by *Kif1c* mRNA localization. **A**. Mass spectrometry of endogenous KIF1C IPs. Samples were prepared in triplicate. Only hits more than than 3-fold enriched over *Kif1c*^*LOF*^ were included. Functional categories were determined using PANTHER and manual curation; many genes fall into more than one category. **B**. Fold change (*Kif1c*^*ΔGA*^ */ Kif1c*^*+/+*^), abundance rank, and brief description of candidate proteins selected for validation. **C-D**. Western blot analysis of co-IP assays with endogenous KIF1C from untagged cells (C) or mCherry-tagged KIF1C (D), using α-KIF1C or α-mCherry antibodies, respectively. Note that endogenous KIF1C is expressed at low levels and is undetectable in input samples under these conditions. **E**. A model for the role of *Kif1c* mRNA localization in regulating protein and cellular behavior: In *Kif1c*^*+/+*^ cells (gray), KIF1C undergoes limited, specific protein-protein interactions and transports cargoes related to mRNA trafficking (e.g. APC) and cell migration (e.g. integrin). In *Kif1c*^*ΔGA*^ cells (purple), KIF1C engages with additional, aberrant interactors. KIF1C is still able to transport APC cargoes, but integrin cargoes are affected through an unclear mechanism (faded α5β1 cargo), which may include the cargo being lost, diminished, or unable to recycle properly. In *Kif1c*^*LOF*^ cells (green), KIF1C is absent and all cargos are affected.

### Loss of Kif1c mRNA localization leads to dysregulated protein-protein interactions

When we plotted the relative change in binding interactions (*Kif1c*^*ΔGA1,2*^ / *Kif1c*^*+/+*^), we noticed a pronounced skew of the volcano plot toward the right, suggesting that most interactions are increased when KIF1C is produced from mis-localized mRNA (Figure 6A). One interpretation of this skew is that proper mRNA localization allows for preferential loading of selected KIF1C interactors that may be co-enriched in protrusions, while KIF1C produced from mis-localized mRNA is loaded more promiscuously in the cell body. To validate these results, we ranked hits based on their absolute abundance in the mass spectrometry data and their fold-change of enrichment in *Kif1c*^*ΔGA1,2*^ compared to *Kif1c*^*+/+*^ cells. From this group, we selected eight proteins that represent a range of differential enrichment values and included a barbed end actin filament capping protein (GSN) and an Arp2/3 complex protein (ARPC2) (Figure 6B). Co-IP assays using an α-KIF1C antibody followed by western blotting confirmed that these protein-protein interactions were increased upon *Kif1c* mRNA mis-localization (Figure 6C). As an independent approach, we also carried out Co-IPs using an α-mCherry antibody in the KIF1C:mCherry tagged cell lines and observed the same results (Figure 6D). Because only one allele of *Kif1c* is tagged, the α-mCherry Co-IPs also allowed us to distinguish non-tagged KIF1C, which co-purified with tagged KIF1C due to dimerization irrespective of mRNA localization. In both untagged and tagged cell lines we observed a reproducible range of enrichment of the validated KIF1C interactors. For example, some proteins (e.g. VIM, CTTN) were modestly enriched in *Kif1c*^*ΔGA1,2*^ cells yet still readily detectable in *Kif1c*^*+/+*^ cells, while other proteins (e.g. GSN, ATL3) were barely detectable in *Kif1c*^*+/+*^ cells. We speculate that the former class of proteins may represent normal interaction partners that become dysregulated in response to mRNA mis-localization, while the latter class may be aberrant interactors that opportunistically bind KIF1C when the mRNA is not translated in the proper context. In conclusion, these data reveal a critical role for *Kif1c* mRNA localization in dictating the specificity of KIF1C protein-protein interactions.

## DISCUSSION

Here, we demonstrate that 1) The role of KIF1C in cell migration and mRNA trafficking are mechanistically separable, 2) KIF1C function is predicated, at least in part, on localization of *Kif1c* mRNA, and that 3) *Kif1c* mRNA localization is required for establishing specificity of protein-protein interactions (see model in Figure 6E). Recently, it was shown that localization of another APC-dependent mRNA, *Rab13*, is required for proper loading of a key interaction partner^6^. The fact that *Rab13* and *Kif1c* are among the first APC-dependent mRNAs to be studied in detail strongly suggests that the role of mRNA localization in binding partner specificity will be a common mechanism in non-neuronal cells. Furthermore, we predict that RAB13 interactions may be more broadly dysregulated than previously appreciated.

Our data show that *Kif1c* mRNA localization regulates cell migration but not mRNA trafficking. We hypothesize these differential sensitivities to mRNA localization could, in some contexts, provide a mechanism for regulating KIF1C activity. For example, increased or decreased localization of the *Kif1c* mRNA would provide a means to divert the pool of KIF1C in order to enhance or suppress migration, as opposed to simply tuning the overall level or activity of the protein which would also affect mRNA trafficking. Furthermore, because this provides a post-transcriptional mechanism for regulating cellular behavior, it would allow for rapid responses to extracellular cues^3,21^. Examination of whether and how the degree of *Kif1c* mRNA localization is regulated is therefore an interesting direction for future investigation.

How *Kif1c* mRNA localization to protrusions modulates protein-protein interactions is unclear. One possibility is that mRNA localization permits only key interaction partners to gain privileged access to nascent proteins in an otherwise crowded cellular milieu. Without sequestration, ubiquitous cellular proteins may out-compete proper interaction partners for binding. The variable effect of mRNA localization on downstream functions may derive from differential sensitivity to changes in binding partners. For example, the role of KIF1C in cell migration requires continual recycling of integrins and is known to be sensitive even to over-expression of wild type protein^17^. As such, subtle changes in binding specificity may be sufficient to inhibit this process. Conversely, while less is known about mRNA trafficking, based on our results we predict the interactions regulating this process will be comparatively more stable and/or robust. Future work will need to address how individual interactors are differentially recruited and loaded, and which are causal for specific phenotypes. We predict that such studies will not only explain the many roles of *Kif1c*, but will also improve our understanding of the functional significance of many other protrusion-localized mRNAs present across non-neuronal cell types.

## MATERIALS AND METHODS

### Cell culture

Established YUMM1.7 and COS1 cell lines were obtained from ATCC. COS1 cells were cultured in DMEM supplemented with 10% FBS and 5% penicillin/streptomycin. YUMM1.7 cells were cultured in DMEM:F12, HEPES with L-glutamine supplemented with 10% FBS, 5% non-essential amino acids and 5% penicillin/streptomycin. The YUMM1.7 cell line was authenticated after receipt using allele specific primers and confirmed to be negative for mycoplasma.

### Fractionation of cells, RNA-seq and qRT-PCR

PET Millicell Hanging Cell Culture Inserts with 1 μm pores sized for 6 well plates were coated on the bottom side with 30 ug/mL fibronectin. Fibronectin was diluted in sterile PBS and inserts were coated for 15 minutes, then aspirated and allowed to dry. 4×10^5^ to 8×10^5^ cells were plated per membrane and placed in an incubator overnight. For fractionation, media was aspirated, cells were rinsed with RNase-free PBS and aspirated, then cell bodies were scraped using a cell scraper and collected into 600 ul RLT buffer from the Qiagen RNeasy kit. Inserts were rinsed with RNase-free PBS and cleaned with a cotton swab three times. The cleaned insert was carefully dislodged, placed inside a new tube with 600 ul RLT buffer and briefly vortexed. After cell bodies and protrusions were collected into RLT for each sample, the standard protocol for RNeasy was followed, including DNase digestion. RNA was eluted with 30 ul RNase-free water. 16 ul of protrusion RNA or 1 ul of cell body RNA was used for making cDNA with Takara PrimeScript RT Master Mix. cDNA was diluted 1:10 and 4 ul was used to carry out qPCR with power SYBR reagents. For qPCR, 1-3 filters were used per sample. For RNA-seq, samples were prepared as above except samples were collected into Qiazol and the Qiagen miRNeasy kit was used to allow for capture of small RNAs. There were two RNA seq replicates, each with 6 inserts and 6-8×10^5 cells per insert, subjected to strand-specific whole transcriptome sequencing at a depth of 25-35 million reads per sample. For qRT-PCR on bulk cellular RNA, RNA was prepared using RNeasy, then quantified and an equal amount from each sample was used to make cDNA; cDNA was diluted 1:20 before use. Stably expressed genes for normalizing qRT-PCR data were identified using NormFinder^22^.

### Lentivirus production, generation of pooled knockout populations and MiSeq

Heterogenous knockout pools were generated using lentiCRISPR_v2 as described previously^23^ using the top two guides for each gene from the Brie sgRNA library^24^. Briefly, guides for candidate genes or two non-targeting guides were cloned into lentiCRISPR_v2 (Addgene #52961). Virus was generated by plating 3.5×10^4^ COS1 cells per well in a 6 well plate and transfecting them on the 2^nd^ day with 1 ug total plasmid per well at a 5:3:2 ratio of lentiCRISPR:psPAX2(Addgene #12260):pMD2.G(Addgene#12259) using FuGENE HD. Viral media was collected after 48 and 72 hours, passed through a 0.45 μm filter and stored as aliquots at -80º. Viral media was diluted 1:1 with YUMM1.7 complete media including 8 ug/mL polybrene and used to transduce YUMM1.7 cells in 12 well plates. Media was changed 24 hours after transduction and 2 ug/mL puromycin selection was started after an additional 24 hours. Cells were selected for 6 days and expanded into 10 cm plates. gDNA for MiSeq was collected on selection day 5 and prepared using Qiagen DNeasy. MiSeq libraries were prepared through two rounds of PCR: 1) gene-specific primers with overhangs; 2) overhang-specific primers for indexing and multiplexing. PCRs from each sample were concentration matched and pooled, run on a 2% agarose gel stained with SybrSafe, gel purified with Qiagen Gel Extraction Kit and submitted for next generation sequencing using MiSeq Reagent Nano Kit v2 at a depth of 1 million reads. Resulting fastq files were analyzed using CRISPResso^25^. Cells were plated for live cell tracking on days 8 and 13 post-selection, and counted for proliferation every 2-3 days throughout.

### Live cell tracking

Live cell tracking of single cells was carried out in 24 well dishes on a heated stage with atmospheric control using a Zeiss AxioObserver Z1 equipped to automatically perform time-lapse live-cell image acquisition. Cells were plated dilutely (∼500 cells per well) in plastic dishes without (Figure 2) or with (Figure 5) fibronectin coating and allowed to acclimate overnight, then imaged using phase contrast and a 10x EC PlnN air objective with NA=0.3 every 3 minutes for 8 hours. Cells were manually tracked using Trackmate in Fiji. Cells that were in view over a 5 hour period were used for analysis. For tracking on fibronectin, cells were excluded if they divided during the tracking window. Tracking data was processed with the chemotaxis tool (Ibidi) then analyzed using Rstudio with a code prepared in house. Briefly, the cumulative distance and speed and net displacement were calculated for every cell at every time point, then the average across cells at the same timepoint was determined for each genotype. For experiments with replicates, the mean of the replicates and the standard error of the mean was plotted for each time point. Cell shape was analyzed by measuring cell features in Fiji of all cells in view during the 50^th^ frame of every movie. Cell length is the longest straight line that can be drawn for a given cell from tip to tip.

### Generation of clonal cell lines with deletions and insertions

To generate clonal cell lines, cells were transfected with px458 which expresses GFP, Cas9 and sgRNA. Guides were selected using the UCSC genome browser -NGG target site function. GFP+ cells were single cell sorted 48 hours after transfection. Clones were expanded and genotyped using primers spanning sgRNA cut sites. For deletions, two sgRNA were used; for indels, one sgRNA was used. The genotype of mutated alleles was determined using Sanger sequencing. For mCherry insertion via homologous recombination one sgRNA targeting just after the stop codon was used and px458 was co-transfected with a plasmid containing linker-mCherry flanked by 1,000+ nucleotide *Kif1c* homology arms in which the PAM site for the sgRNA was altered to inhibit recutting after repair. Transfected cells were first sorted as GFP+ pools after 48 hours, then single cell sorted for mCherry+ cells 6 days later. The tagged and untagged alleles were analyzed using Sanger sequencing. sgRNA sequences are in a Supplementary File 1.

### Live cell fluorescent imaging and analysis

Live cell imaging of fluorescent cells was carried out in Nunc Lab-Tek II 8 well chambered coverglass in Live Cell Imaging Solution (Invitrogen) using a 40x EC PlnN oil objective with NA=1.3 on a Zeiss AxioObserver Z1 with an AxioCam MRm monochrome digital camera. Coverglass were coated with 30 ug/mL fibronectin diluted in PBS for 15 minutes, aspirated and allowed to dry. mCherry-tagged cells and an untagged parental cell line were plated dilutely and allowed to adhere for 2 hours. Cells were rinsed and then imaged in pre-warmed imaging media. Six slices, 0.7 μm apart, with 1500 ms exposure time were captured for each cell using an HXP 120C light source and Filter Set 63HE, as well as a bright field image. Cells were analyzed using Fiji as follows: Cell outlines were manually drawn. Cell bodies vs protrusions were distinguished using white light.

The cell body was defined by edge of the rough matter surrounding the nucleus, which coincided with flexion points in the cell membrane near protrusions. Only the slice in which the base of the cell was sharply in focus was quantified. Only cells that unambiguously did not overlap with other cells or debris were used. Protrusion enrichment was calculated as follows:

1. The mean fluorescence per unit area for protrusions and cell bodies was calculated for every cell, including negative controls. 1) The integrated density and area of the whole cell and the cell body were measured. 2) Protrusion integrated density = whole cell integrated density - cell body integrated density. Protrusion area = whole cell area - cell body area. 3) Mean Fluorescence (cell body or protrusions) = integrated density / area.

2. The average mean fluorescence in protrusions and cell bodies due to background autofluorescence was calculated by averaging the measurements from 10+ negative control cells for each experiment.

3. The average protrusion-specific and cell body-specific background (quantified in step 2) was subtracted from protrusions and cell bodies of tagged cells.

4. The corrected protrusion and cell body fluorescence values were used to calculate fluorescence in protrusions/cell bodies.

Whole cell fluorescence of tagged cell lines was measured using a BD Melody FACS and FlowJo software. 12,000-16,000 tagged single cells per cell line were measured using the PE-CF594 laser.

### Western blot and Co-immunoprecipitations

Bulk protein was collected from confluent cells grown in a 6 well plate. Cells were rinsed with ice cold PBS then scraped in 80 ul of cold RIPA buffer (50 mM Tris-HCL, pH7.4, 1% NP-40, 0.5% Na-deoxycholate, 0.1% SDS, 150 mM NaCl) with 1x Complete, EDTA-Free Protease Inhibitor (Sigma). Lysates were agitated at 4º for 20 minutes, pelleted, and the supernatant transferred to a new tube. For Co-IPs, 4×10^5^ – 1.5×10^6^ cells were plated 2-4 days before collection in 15 cm dishes such that they were 90% confluent on the day of collection, with 2 dishes per sample. Samples were prepared in the cold room by rinsing briefly with ice cold PBS, aspirating the PBS and scraping the cells into a 1.5 mL tube using residual PBS. Cells were immediately spun for 1 min x 1,000 g, aspirated, and 300 ul Thermo Cell Lysis Buffer (NN0011) with 2x Complete Protease Inhibitor and 2x PhosSTOP was added. Lysates were resuspended with gentle pipetting, then rotated for at least 30 minutes at 4º, pelleted 10 min x 14,000g and supernatants moved to a new tube. Lysates were precleared with 50 ul washed Protein G Dynabeads for 30 minutes, then beads were discarded and lysate concentration was measured using Pierce BCA Protein Assay Kit and the volume and concentrations for each genotype (e.g. +/+, ΔGA and LOF) were matched using extra lysis buffer as needed. For mass spec, 4.5-5 mg total protein were used for each replicate with 4 ug of KIF1C antibody (Bethyl Kif1c/LTXS1 antibody, A301-072A). For western blots, 3-4 mg total protein and 2 ug of KIF1C antibody or 10 ug mCherry antibody (Invitrogen mCherry Monoclonal Antibody (16D7)) were used. Antibody was added directly to lysates and the mixture was rotated at room temperature for 90 minutes, then 75 ul of rinsed Protein G Dynabeads were added and rotated for an additional 30 minutes at room temperature. Unbound proteins were removed by washing 3x with wash buffer (50 mM Tris, pH 7.4; 150 mM NaCl; 0.02% NP-40; 1x PhosSTOP and 1x Complete Protease Inhibitor), then transferring the beads to a fresh tube and eluting in 18ul 1x Laemmli buffer (Boston bioproducts BP-111R). All of the eluate was run on NuPAGE™ 4-12% Bis-Tris Protein Gels, 1.5 mm, 15-well. For Mass spec, the proteins were resolved for 100 mm, then labeled with SimplyBlue SafeStain and carefully excised and submitted for analysis on a Lumos mass spectrometer. Data was analyzed with Proteome Discoverer 2.4 and searched using the mouse protein database from UniProt. Proteins that were identified by a single peptide or identified but not quantified were omitted from analysis. For western blotting, SDS Page was run on NuPAGE™ 4-12% Bis-Tris Protein Gels until the 20 kDa band of the protein ladder ran off the gel, then the proteins were transferred to nitrocellulose membranes, blocked with milk and blotted 1:1000 with designated antibodies: From Proteintech: TPM1/28477-1-AP; ARPC2/15058-1-AP; CTTN/11381-1-AP; GSN/11644-2-AP; RAI14/17507-1-AP; FLII/67039-1-Ig; ATL3/16921-1-AP. From Santa Cruz: VIM/sc-6260. From Sigma: TUBULIN/T5168. Secondaries were used at 1:10,000 dilution: IRDye 800CW Donkey anti-Rabbit or anti-Mouse IgG; IRDye® 680LT Goat anti-Rabbit or anti-Mouse. For ATL3 and VIM, a light chain specific secondary was used: IgG Fraction Monoclonal Mouse Anti-Rabbit IgG, light chain specific 790. Blots were imaged on a Li-cor Odyssey Clx imaging system and analyzed with ImageStudio.

## SUPPLEMENTARY DATA

Supplementary file 1:

- Guide sequences used for Lenticrispr_v2
- Guide sequences used for generating clonal cell lines
- Genotyping primers for analyzing CRISPR alleles
- Primers for qPCR
- Primers for MiSeq

Supplementary table 2:

- RNA Seq results

Supplementary table 3:

- Mass spec results

## ACKNOWLEDGEMENTS

We would like to thank Dr. Khuloud Jaqaman for her advice on quantitative image analysis, Dr. Adam Norris for helpful comments on the manuscript and the UT Southwestern Proteomics, Next Generation Sequencing and Flow Cytometry cores for their assistance in mass spectroscopy, high-throughput sequencing and fluorescence activated cell sorting, respectively. Figures created in part with BioRender.com.

